# Event Segmentation and Linguistic Granularity in Direct/Indirect Causation: Unraveling the Mind-Language Interface

**DOI:** 10.64898/2026.07.01.733328

**Authors:** Mengmin Xu, Yan Ren

## Abstract

Building upon foundational psychological theories of event segmentation, this study addresses the limitation of overreliance on temporal boundaries as the primary segmentation criterion. Drawing on two experiments of direct and indirect causation in Mandarin Chinese, this study demonstrates how cognitive segmentation granularity and semantic integration jointly shape syntactic encoding. Results reveal distinct event encoding patterns for direct and indirect causation: coarse-grained segmentation leads to compact syntactic structures (e.g., verb-resultatives), while fine-grained segmentation yields varied multi-clausal expressions. Chinese speakers update event models via prediction errors of intentionality and protagonists, and tend to establish event boundaries at goal-relevant action endpoints when construing causal chains. These conceptual dimensions exert a modulating influence on both event segmentation and semantic integration. We propose a triad model integrating event segmentation, semantic integration, and linguistic specificity, providing a unified framework for elucidating the mind-language interface in conceptual construction and event coding of causation.

## 1. Introduction

Events, cognitively conceived and temporally bounded segments of experience (Radvansky & Zacks 2014; Zacks & Tversky 2001; Zacks et al. 2007; Zwaan 2016), constitute fundamental units through which individuals perceive and structure the world. Through cognitive processing, individuals parse continuous streams of activity into discrete events and sub-events, thereby employing linguistic structures to interpret them. This mental construction aligns with Slobin’s (1996) hypothesis, which posits that the world itself does not represent events; rather, human experiences of events are constructed based on linguistic frameworks where events are encoded within language. The mechanism underpinning this construction is *event segmentation*: the cognitive process of demarcating and interpreting coherent units from ongoing activity streams (Deng & Li 2023; Gerwien & von Stutterheim 2018; Newston 1973; Radvansky 2017; van Staden & Narasimhan 2013; Wang et al. 2023; Wolff et al. 2009; Zacks 2020; Zacks & Tversky 2013; Zacks & Swallow 2007). Consequently, event segmentation and linguistic representation are intrinsically linked, forming complementary processes. Extending this line of inquiry, Bohnemeyer et al. (2007, 2010) demonstrate that a single causal event can be semantically construed in multiple ways. For instance:

1. a. Sally hit the vase. It fell and broke.

Sally knocked over the vase. It broke.
The vase broke because Sally knocked it over.
Sally broke the vase. (Bohnemeyer et al. 2007: 496)

Analyses of the vase-breaking event reveal distinct segmentations: three sub-events in (1a), two in (1b), clausal embedding for cause in (1c), and a highly abstract single-chain event in (1d). This spectrum of representations directly demonstrates the interplay between event segmentation and linguistic structure.

Theoretical accounts of what “anchors” event segmentation have traditionally fallen into two distinct camps. Linguistic approaches contrast with psychological ones. The first, primarily from linguistics, focuses on symbolic units. Proposals within this camp advocate for syntactic units like verbs and clauses (Pawley 1987) or intonation (Givón 1991) as key segmentation criteria. A notable contribution is Bohnemeyer et al.’s (2007) macro-event property, which serves as a formal and explicit syntactic criterion for determining how sub-events are packaged into coherent macro-events in language. Specifically, this principle centers on the scope of temporal operators: if a single temporal operator (e.g., adverbs of time) can quantify over a sequence of contiguous sub-events, those sub-events are deemed to constitute a single macro-event. In this way, the macro-event property provides a concrete methodological tool for measuring event density—i.e., the degree to which multiple sub-events are integrated into a single syntactic unit—by examining the coverage of temporal operators over event sequences. A limitation of this model, however, is its sole dependence on temporal features, which overlooks other semantic parameters and cross-level granularity differences that also play crucial roles in event segmentation and syntactic packaging. Secondly, psychological perspectives ground segmentation in cognitive processes, emphasizing the role of perceptual event features and action goals. Studies show that observers segment events when perceptual features contradict expectations (Zacks & Swallow 2007; Zwann & Radvansky 1998). Concurrently, action goals serve as a key cognitive mechanism for segmentation, whereby individuals infer relationships between (sub)events by tracking an agent’s goals (Zacks & Tversky 2001, 2013). Bridging this theoretical divide, the present study integrates evidence from both symbolic units and cognitive processes to explore the interface between conceptual event segmentation and its linguistic realization.

Research on linguistic event segmentation remains in a nascent stage. Seminal work by Davidson (1969) laid the groundwork by advocating for causation as a basis for segmentation. Building on this, Wolff (2003) proposed the “no-intervening-cause criterion”, positing that direct causal chains — lacking an intervening agency or where an intermediary serves as a tool — are encoded as single macro-events, whereas causally mediated events are segmented into two distinct units (see Figure 1). Parallelly, Bohnemeyer et al. (2007, 2010) investigated event segmentation and semantic density in motion events and change-of-state events using the macro-event property, focusing on cross-linguistic typology. Meanwhile, Liu (2016) highlighted that boundary assignment underpins the establishment of linguistic structure and serves as a cognitive motivation for lexicalization. Despite these valuable contributions, the finer granularity of event segmentation has received scant attention. Crucially, the criteria governing event boundary establishment and their consequent impact on syntactic organization remain unexamined.

**Fig 1.**
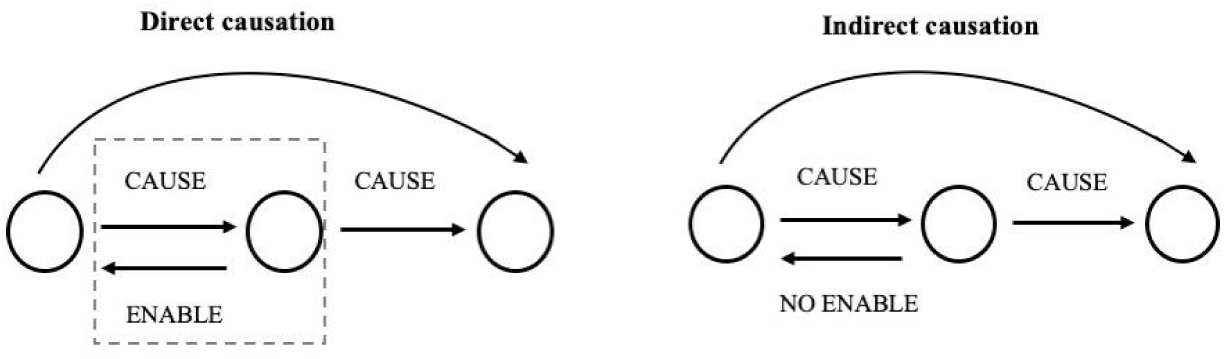
Direct and indirect causation specified by the no-intervening-cause criterion (Wolff 2003)

Against this backdrop, this study examines how different causal chains are segmented and linguistically encoded in Chinese. Causal relationships, being fundamental to human cognition, are prototypically conceptualized in language as sequences of interconnected events or sub-events (Talmy 2000: 271). This conceptualization makes them an ideal domain for investigating event segmentation, as they integrate physical changes with intentional action. However, the cognitive mechanisms that govern the segmentation differences and the conceptual construction of direct and indirect causation remain underexplored. Through two sets of comparative experiments, this research addresses three questions concerning how event parameters influence segmentation and syntactic organization: (1) What segmentation strategies do native Chinese speakers adopt for direct and indirect causal events? (2) How are these distinct causal events mapped onto linguistic representations in Chinese? (3) To what extent do event segmentation and relevant parameters regulate the syntactic organization of causal expressions in Chinese?

## 2. Segmentation, Sub-event Integration and Clausal Union

### 2.1 Event segmentation in cognition

According to Zacks (2004), the human cognitive system parses the flow of experience into discrete, meaningful units. This process involves the dynamic mental representation that segment ongoing activity into coherent parts, thereby constructing event contours and enabling understanding of internal event structure. The demarcation between these segments is known as an event boundary or “breakpoint” (Krebs et al. 2025; Zacks & Tversky 2001; 2013). The perception of a structured event gestalt relies on these boundaries, which turn continuous activity into a cohesive whole composed of bounded internal units. As boundary assignment is central to segmentation, it directly governs the architecture of conceptual representations.

Critically, event segmentation occurs at multiple levels of granularity, commonly categorized as coarse-grained (CES) or fine-grained (FES) (Zacks & Swallow 2007). This process is hierarchical: coarse-grained segmentation identifies abstract, overarching events, while fine-grained segmentation delineates their constituent sub-events (Keven 2016). For example, the high-level event “going to buy a book” comprises finer-grained actions such as “walking to the bookshelf”, “selecting a book”, and “paying”. Notably, event segmentation involves not only information extraction but also goal-directed decomposition, breaking down overarching activities into constituent sub-actions (Zacks & Tversky 2001).

A well-established theoretical account of event segmentation is the event-indexing model (Zwann & Radvansky 1998). This model has provided valuable insights for the field by identifying five situational dimensions that trigger event boundaries: time, space, protagonists & objects, causation, and intentionality. It argues that segmentation boundaries emerge at points of discontinuity along these dimensions, and the likelihood of perceived boundaries increases with the number of changing dimensions. A major strength of the model is that it adopts a multi-dimensional framework in boundary assignment, which goes beyond accounts centered only on perceptual and spatio-temporal cues. In doing so, it highlights the role of conceptual factors in event segmentation, providing an important foundation for the present research. However, the event-indexing model also exhibits certain limitations that motivate the present study. First, although the model identifies five dimensions that trigger boundaries, it does not specify how these dimensions interact or whether some dimensions take priority over others in guiding segmentation. Second, the model focuses primarily on where event boundaries arise during cognitive processing, rather than systematically explaining how these factors influence the further integration, structural packaging, and syntactic mapping of event units during language production. As a result, it does not connect cognitive segmentation boundaries with linguistic iconicity—particularly in languages like Mandarin Chinese, where discourse and pragmatic factors may weigh differently.

These gaps leave open the question of how conceptual (sub)event units, once segmented, are integrated and expressed in language. These limitations underscore the need for a framework that systematically links event segmentation boundaries and conceptual construal from psychology onto syntactic structure. The present study addresses this need by combining insights from the event-indexing model with theories of semantic integration and clausal union, as elaborated in the following sections.

### 2.2 Semantic integration and specificity

As a salient and universal event frame, the linguistic representation of a causal chain is not solely determined by underlying conceptual segments but is equally governed by the specific way in which a conceptualizer construes that content (Langacker 2008: 77). Construal refers to the cognitive ability to structure a situation through operations such as setting a conceptual scope, imposing profiling (foreground/background organization), and adjusting specificity (granularity) (Langacker 1987: 138).

The cognitive process underlying event segmentation and semantic integration of causal chains can be explained by Talmy’s (2000) principle of the “windowing of attention”. This principle provides a mechanism for conceptual highlighting, whereby language structures foreground certain portions of an event frame through explicit mention (windowing), while backgrounding others through omission (gapping). Applied to causal events, the sequential structure of a causal chain offers a cognitive substrate for this process. This mechanism allows speakers to represent an extended causal chain by selectively directing attention to specific, salient sub-events. The choice of which causal sub-events to window or gap directly governs semantic integration of the segments between an initiating action and its ultimate outcome. The degree of linguistic granularity is thus a direct reflection of event segmentation and semantic integration of the causal sequence. For causal chains, Talmy (2000: 257) formalizes event segmentation with an initiatory intentional agent through the following event frame:

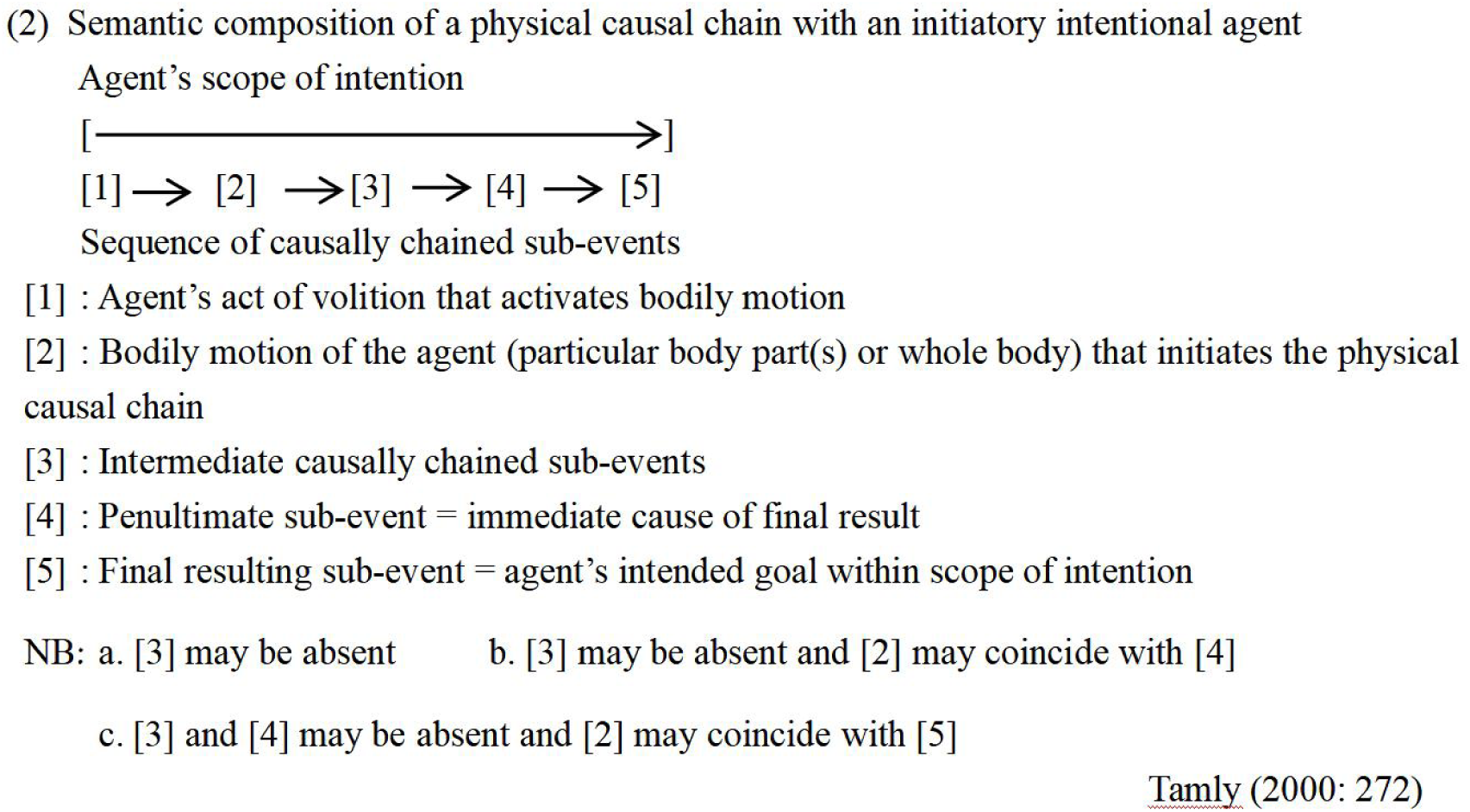

The conceptualization of a causal chain involves segmenting a continuous flow of experience into relatively discrete sub-event “packets”, where the sense of causality itself may reside primarily in the boundaries between a sub-event and its successor (Talmy 2000). Building on this framework, the present study proposes that direct and indirect causal chains are semantic-conceptual complexes composed of multiple sub-events. Their conceptual differentiation and linguistic construction are driven by distinct cognitive operations pertaining to event segmentation, the degree of specificity, and sub-event integration. Through these mechanisms, the mental representations of direct and indirect causation are shaped and mapped onto language.

### 2.3 Clausal union in language

The systematic isomorphism between semantic and syntactic structure offers a cross-linguistically robust paradigm of syntactic iconicity (Haiman 1983; Givón 1991; Papafragou & Grigoroglou 2019). This iconic interface is particularly critical for representing causal events, where the semantic dimension of event segmentation and integration (the “semantic bond”) maps directly onto the syntactic dimension of clause union (Givón 2001). Consequently, the cognitive distinction in the degree of event segmentation and conceptual integration should be systematically reflected in the tightness of syntactic packaging. This cognitive-syntactic interface is governed by the key principle whereby the strength of the semantic bond between two events predicts the degree to which their clauses are syntactically integrated into a complex unit (Givón 2001, 2009). This principle defines that syntactic complexity (clause union) is a direct reflection of cognitive-semantic complexity (event integration).

The linguistic representation of a causal event involves a complex internal structure where sub-events are organized into a systematic whole based on cognitive connectivity (Givón 2009; see Figure 2). This process of structuring is guided by Gestalt principles, which drive the segmentation of the causal continuum into discrete event and sub-event units. Givón identifies six semantic-cognitive dimensions that are pivotal for event integration and clausal union. These include dimensions related to core *eventhood*: co-temporality (temporal integration), spatial contiguity (spatial integration), and co-reference (referential integration); and dimensions related to *agentivity*: intentionality, control, and coercive power. Notably, Givón’s framework for event integration and the event-indexing model (Zwann & Radvansky 1998), while developed independently, converge conceptually. The dimensions of temporal, spatial and referential integration in Givón’s system directly elaborate on the “time” “space” and “protagonists & objects” dimensions in the event-indexing model. Similarly, the *agentivity* cluster in Givón’s theory deepens the interpretation of “protagonists”, “causation” and “intentionality” dimensions. Crucially, these two frameworks are complementary. The event-indexing model helps identify where segmentation boundaries arise, while Givón’s principle of event integration clarifies how these segmented units are conceptually integrated and linguistically structured. In this sense, the event-indexing model provides a robust account of boundary detection, wherea Givón’s framework offers a theory of syntactic iconicity that connects semantic integration directly to clausal structure.

**Fig 2.**
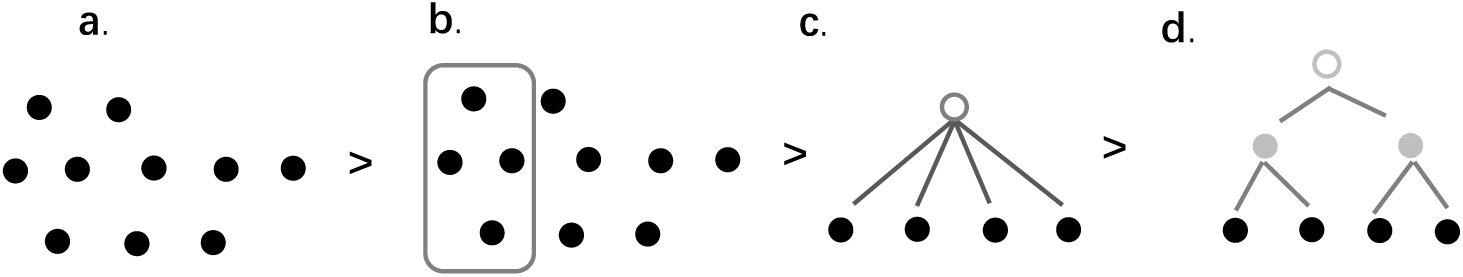
Portion excerpting and hierarchical systems of organized structures (based on GivÓn 2009)

Against this theoretical background, the present study proposes and empirically tests an integrated framework that combines the event-indexing model’s account of segmentation boundaries with semantic integration and Givón’s theory of clause union. By synthesizing these approaches, we aim to provide a more comprehensive account that links cognitive segmentation, conceptual integration, and syntactic encoding—a gap that existing individual models have not fully address, particularly for causal event representation. Building on prior work, this study offers several distinctive features. First, we investigate how conceptual factors—causal structure (direct vs. indirect) and intentionality (intended vs. unintended)—shape segmentation granularity and its linguistic expression, moving beyond previous work focused primarily on temporal or spatial boundaries. Second, we examine segmentation and linguistic encoding together within a single experimental paradigm, tracing the cognitive pathway from conceptual parsing to syntactic packaging. Third, focusing on Mandarin Chinese — rich in verb-resultatives and periphrastic causatives—we provide cross-linguistic evidence extending findings from English and other Indo-European languages.

From this integrated framework, we derive our central hypothesis: the perceived directness of causal relationships, modulated by intentionality and mediation, determines the granularity of cognitive event segmentation and the degree of semantic integration; these cognitive processes are iconically reflected in Mandarin clausal structure, with systematic mapping between segmentation granularity and clausal representation complexity. To this end, the following experiments are designed to answer two core questions: First, how does the cognitive segmentation of direct and indirect causal chains vary in granularity? Second, how are these variations mapped onto specific patterns of clausal union in Chinese causal expression?

## 3. Research Methodology

### 3.1 Experimental design

This study employed an experimental paradigm to investigate the segmentation and linguistic representation of direct versus indirect causal events in Chinese. Adopting the established methodology of Wolff (2003), we utilized short video clips (averaging 10.5 seconds in duration) depicting fixed spatial scenes. This design served to minimize variation in temporal and spatial parameters, thereby allowing for a focused examination of how event protagonists, action goals, and causal relationships influence segmentation and boundary assignment. Two sets of comparative experiments were designed to contrast causal events with and without intervening agencies, as well as those with and without explicit action goals. The experimental materials and design are available at: https://osf.io/fwz8t/.

#### Experiment 1: Event Segmentation and Syntactic Encoding of Mediated and Unmediated Causal Chains

Experiment 1 investigated the influence of event protagonists on segmentation by comparing direct causal events (without intervening causes) and indirect causal events (with intervening agencies). In a direct causal event, a single agent brings about a change either directly or by using a tool (*e.g. The girl smashed the cup*). In contrast, an indirect causal event involves a chain of agency, wherein an initial agent triggers an action in an intervening agency, which then causes the final change (*e.g. The man asked the other man to smash the cup*).

The stimuli comprised 20 video clips (Clips 1-20), sourced from video libraries recorded by the Semantic Typology Lab at the University at Buffalo (https://ubstlab.wordpress.com) and supplemented by our own recordings. Clips 1-10 depicted direct causal chains without intervening agencies, whereas Clips 11-20 depicted indirect causal chains involving an intervening agency. The depicted actions included tearing paper, swinging a swing, throwing a ball, smashing a cup, breaking an egg, turning off a desk lamp, closing a door, blowing out a candle, cutting paper with scissors, and breaking a plate.

#### Experiment 2: Event Segmentation and Syntactic Encoding of Intended and Unintended Causal Chains

Experiment 2 investigated the influence of action goals on event segmentation by comparing intended causal events (with explicit action goals) and unintended causal events (without action goals). In an intended causal event, an agent purposefully brings about a change, either directly or instrumentally (*e.g. A girl broke the vase with a ball*). Conversely, in an unintended causal event, an author (see Talmy 2000: 514) causes a change accidentally while performing an unrelated action, with no goal directed toward the outcome (*e.g. A girl accidentally broke a vase while playing with a ball*).

The experiment utilized 20 video clips (Clips 21–40), sourced from animations developed by the Psychology and Language Laboratory at Emory University (http://psychology.emory.edu/cognition/wolff/animations.html#wolff2009) and supplemented by our own recordings. Clips 21-30 depicted intended causal chains with clear action goals, while Clips 31-40 depicted unintended causal chains where the outcome occurred accidentally. The stimuli encompassed a range of actions, including breaking a vase with a ball, extinguishing a candle with a water gun, collapsing a card tower with a playing card, shaking a plate with a hammer, smashing a cup by hand, chopping wood with an axe, hitting a ball with a racket, pushing a swing, toppling a paper cup tower, and hitting a swing with a racket.

### 3.2 Participants

Participant recruitment started on 05/03/2024 and ended on 26/03/2024. A total of 30 native Chinese speakers (aged 18-32) were recruited through random sampling from universities and communities in Beijing. The sample was gender-balanced (15 females and 15 males), and all participants were highly educated (holding at least a bachelor’s degree). None of the participants had received formal education or training in linguistics (e.g., courses related to syntax, semantics, or event cognition) to avoid potential biases in task performance. All participants voluntarily consented to participate in the experiment and signed an informed consent form prior to the start. Each participant was assigned a unique identifier from S1 to S30.

### 3.3 Procedures

The experiment began with participants completing a basic demographic questionnaire (e.g., name, gender, age, place of origin, and educational background). The entire interview process adhered to the principles of non-coercion and non-prompting to ensure the naturalness of the spoken Mandarin data collected. A one-on-one experimental session followed, in which participants viewed a series of video clips. The experiment was conducted in a quiet room, with stimuli presented on a laptop computer. After reading a brief instruction screen, participants were asked to remain close to both the screen and an audio recorder. Participants were firstly given the plain instruction in Mandarin Chinese to guide their simple, intuitive judgments based on natural cognitive segmentation of the visual stimulus. To address this, we provided a brief example to clarify that some sequences could reasonably be perceived as one event or multiple separate sub-events in the experimental instruction. Following this non-verbal task, participants were asked the neutral open-ended question “What happened?” to elicit their natural perception and description of the scene. Before proceeding to the formal experimental clips, participants first completed two test video clips to familiarize themselves with the task. The experimenter confirmed that participants fully understood the task, including the sequence of operations and the requirements of each step.

“您将观看 40 段短视频。每段视频结束后, 请依次完成两个任务:

1. **事件切分**: 请判断视频中有几个独立的(子)事件。
2. 语言表征: 请自然、流利地口头描述视频中“发生了什么(”。*注意: 如果您对内容不确定, 可以回放。*)”

You will watch 40 short video clips. After each clip, please complete the two tasks in sequence:

1. **Event Segmentation**: Please judge how many separate (sub) events there are in the video clip.
2. **Linguistic representation**: Please verbally describe what happened in the video naturally and fluently. (Note: If you are unsure about the content, we can replay the clip.)

The formal video clips were played in sequence. The next clip was presented only after the participant had completed both the event segmentation and the oral description. Each participant viewed and described the full set of 40 video clips before the next participant began. To control for order effects, the presentation order of the video clips was counterbalanced across participants using a Latin square design and randomly assigned within each participant’ s experimental session. This randomization and counterbalancing ensured that no systematic order bias affected participants’ judgments. For each video, participants performed the non-verbal and verbal tasks in sequence. In the event segmentation task, participants judged the number of discrete events in the video and gave a numerical response. In the linguistic representation task, participants verbally described the observed scene. This task followed the segmentation task to ensure that the verbal description did not bias the segmentation judgment.

The entire experiment was conducted in Mandarin Chinese and audio-recorded. The recordings were later transcribed verbatim, and two independent coders (blind to the experimental hypotheses) annotated and analyzed the participants’ event segmentation judgments and linguistic representations. In terms of the analysis of participants’ responses: After transcription, the two independent coders first annotated the number of sub-events reported by each participant and the number of clauses in their verbal descriptions. Inter-coder reliability was calculated using Cohen’s kappa, yielding a kappa value of 0.92, indicating high inter-coder consistency. Discrepancies between coders were resolved through discussion until a consensus was reached. For the linguistic representation data, the coders further annotated causal expressions (e.g., the use of causal connectives, verb types). Descriptive statistics and inferential statistical analyses (e.g., *t*-tests) were then conducted to examine differences in event segmentation and clausal numbers across different causal types. This process yielded a total of 1,200 causal expressions (30 participants × 40 instances per participant) of contemporary spoken Mandarin describing direct and indirect causal events.

Regarding the consistency between the number of sub-events in participants’ verbal descriptions and the number of sub-events they reported. The matches and mismatches between these two measures were both recorded and analyzed as an essential part of the dataset. A mismatch was defined as a divergence between the number of sub-events explicitly encoded in the verbal description and the number of sub-events participants explicitly reported. All mismatches were documented in detail, and their distribution across different types of causal chains was analyzed to explore the potential relationship among cognitive segmentation, semantic integration, and linguistic expression.

## 4. Results

Through systematic data classification and statistical analysis based on Xing’s (2004) criteria for Chinese causal sentences, we examined the segmentation and linguistic encoding of direct and indirect causation. The principal findings are summarized as follows:

### Finding 1: Segmentation patterns of direct and indirect causal chains

The analysis revealed distinct segmentation patterns corresponding to event type. In unmediated direct causal events, sequences were most commonly perceived as single-chain events (79.7%), followed by two-chain (19.3%) and three-chain (1%) patterns. Conversely, mediated indirect causal events exhibited greater segmentation diversity, with two-chain events being most prevalent (70%), followed by three-chain (24.6%), single-chain (3.7%), and four-chain (1.7%) patterns (see Figure 3).

**Fig 3.**
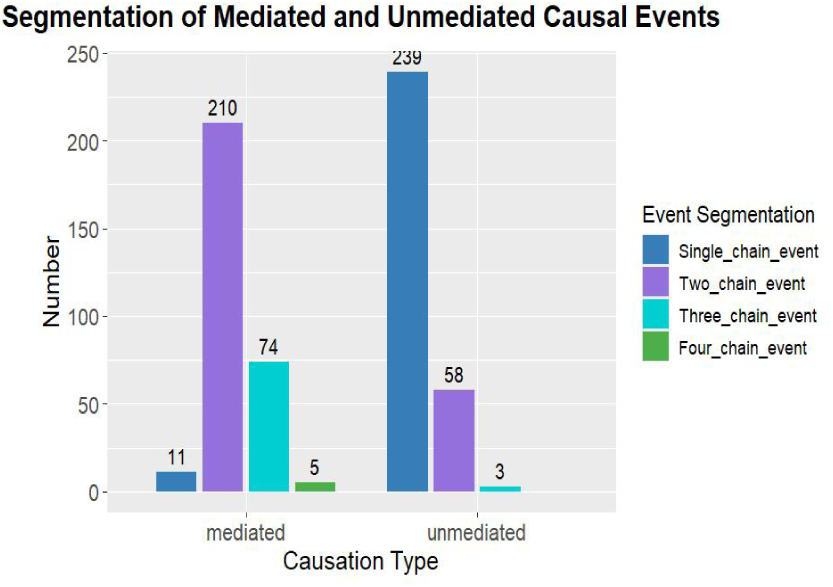
Event segmentation of mediated intended & unmediated causal chains.

A similar distinction was observed in goal-oriented contexts. Intended causal events were predominantly segmented as single-chain (74.6%) or two-chain (20.7%) events, with three-chain patterns being rare (4.7%). In contrast, unintended causal events are most frequently parsed as two-chain (42%) or three-chain (41%) patterns, with single-chain (7%) and four-chain (10%) segmentation being less common (see Figure 4).

**Fig 4.**
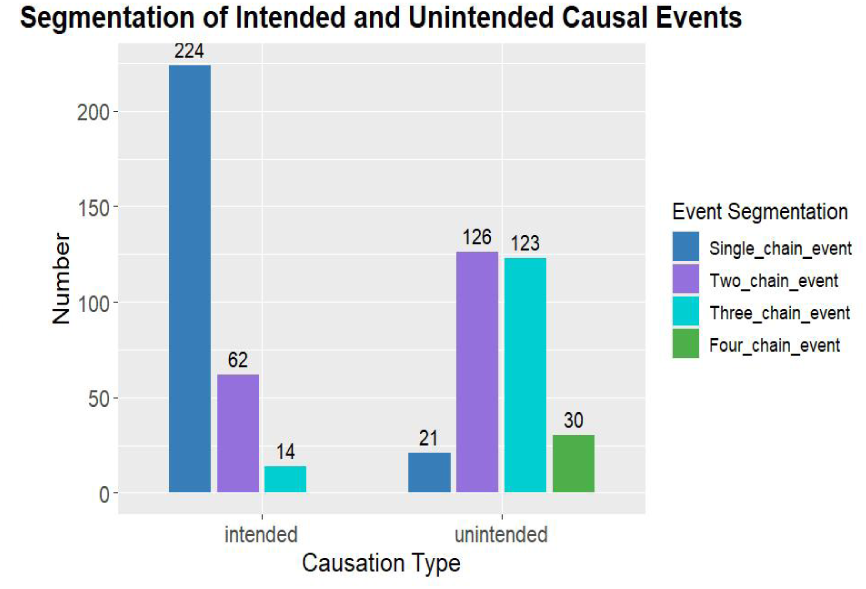
Event segmentation of & unintended causal chains.

### Finding 2: Conceptual construction of direct and indirect causal chains

The syntactic packaging of (non-)mediated causal events varied systematically with the presence of an intervening agency. Descriptions of direct causation heavily favored single-clause constructions (84%), primarily realized as verb-complement predicates / resultative verb constructions (RVCs) (56.7%) that integrate macro-events into a compact syntactic unit. Complex sentences for direct causation (15%) included: 1) causative coordinate clauses (“cause, effect”) linked by conjunctions such as then or thus; 2) causative subordinate clauses (“cause, cause+effect”); and 3) causative divergence clauses (“prerequisite, cause+effect”). Three-clause structures typically extended these conjunctive patterns (1%).

**Fig 5.**
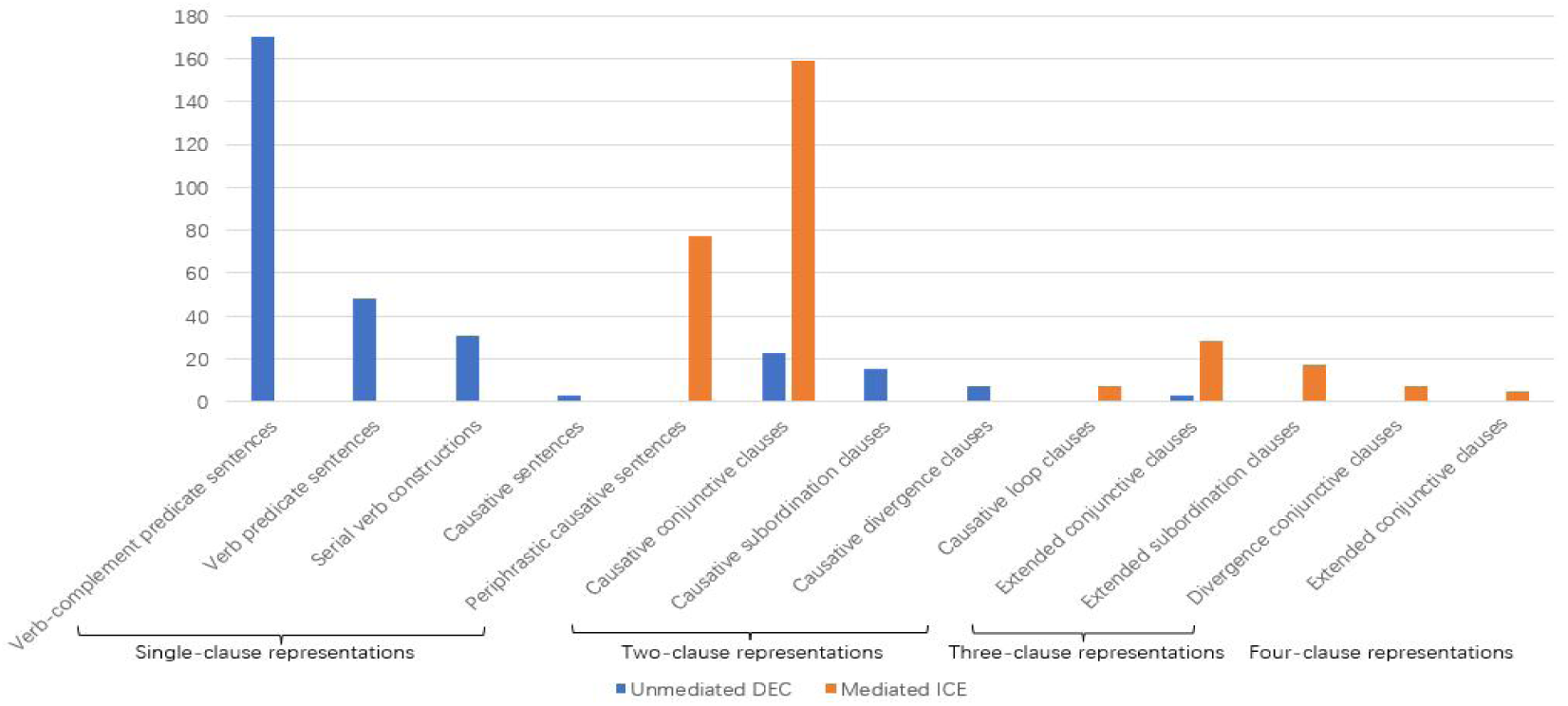
Syntactic encoding of direct and indirect causal chains with & without intervening agencies.

For indirect causation, single-clause descriptions primarily featured periphrastic verbs that compactly integrated “cause1+cause2+effect” (25.7%). At the complex-sentence level, causative coordinate clauses were dominant (53%), encoding relations like “cause, effect” or “cause1, cause2+effect”, occasionally forming causative loop structures (“effect+cause, effect”). Multi-clause structures (3-4 clauses) (19%) involved extended coordinate, subordinate, and mixed divergence-inclusive types.

The absence and presence of an action goal also significantly influenced syntactic form in (non-)intentional causal chains. For intentional causation, single-clause descriptions were frequent (63%), with serial verb constructions being the predominant type (35.3%). Complex sentences included coordinate, subordinate, and divergence forms (32.4%), while three-clause structures mainly featured extended coordinate (“cause, cause, effect”) or divergent-subordinate (“prerequisite, cause, cause+effect”) patterns (4.6%).

**Fig 6.**
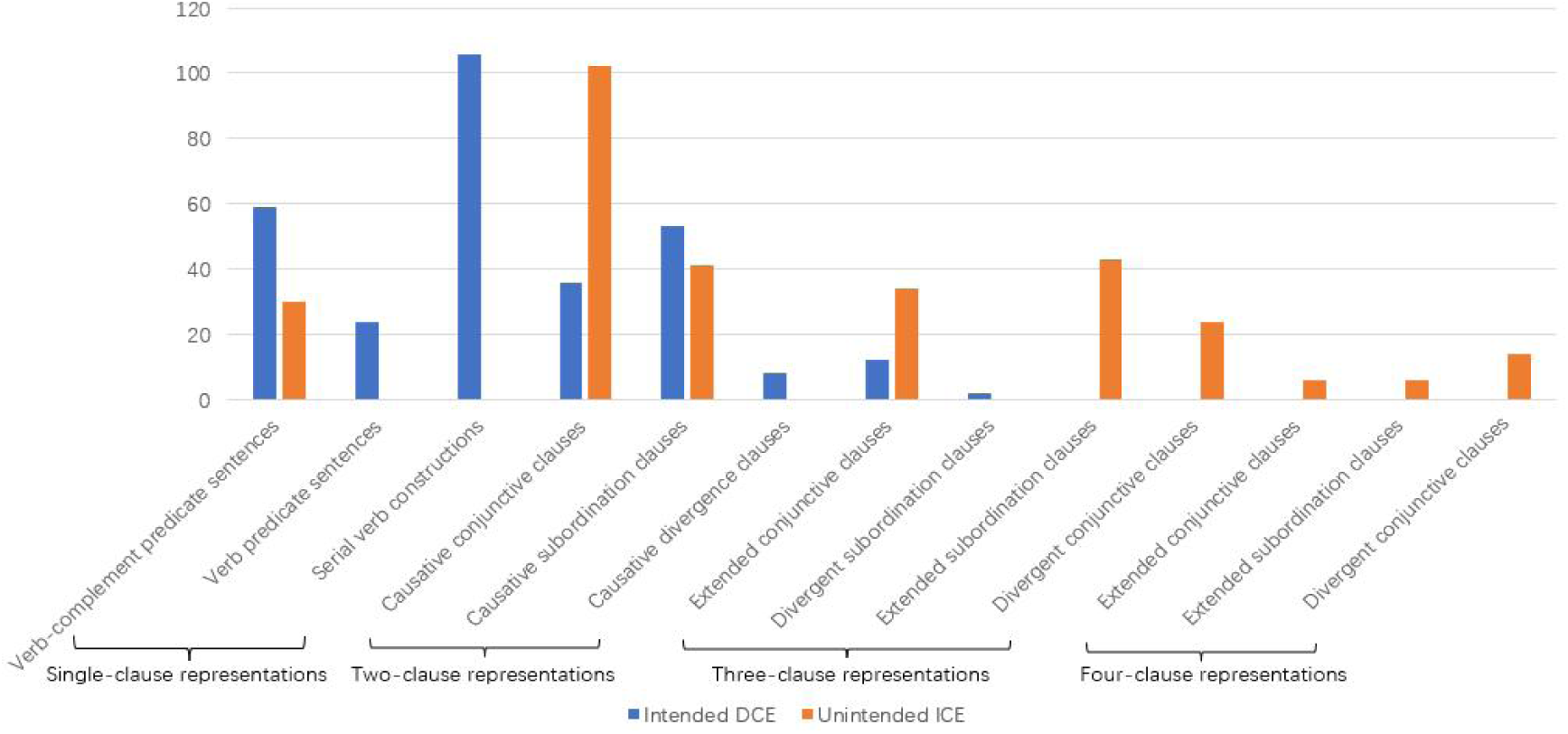
Syntactic encoding of direct and indirect causal chains with & without intended agents.

In descriptions of unintentional causation, single-clause forms were less common (10%) and typically featured verb-complement predicates that packaged “cause+effect” into a single unit, often modified by adverbs like accidentally to underscore the lack of intent. At the complex-sentence level, coordinate clauses were most frequent (34%), sometimes accompanied by subordinate clauses (13.7%). These were often linked by: 1) adversative conjunctions (e.g., but, however) highlighting result unexpectedness, or 2) asymdetic sequencing. Multi-clause structures (3-4 clauses) exhibited looser, extended forms of subordinate, coordinate, and divergent-conjunctive types (42.3%).

### Finding 3: The relationship between event segmentation and clause representation

The descriptive statistics and paired *t*-test results for event segmentation and clause representation across four types of causal chains indicates that the matching rate between event segmentation and clausal representation was as high as 92.5% on average for direct causal chains, and 80.4% on average for indirect causal chains (see Table 1). For direct causal chains, the mismatching cases (7.5%) exhibited fine-grained segmentation (two-chain events) but were conceptually packaged into single-clause representations, indicating semantic integration and clause union processes.

**Table 1.**
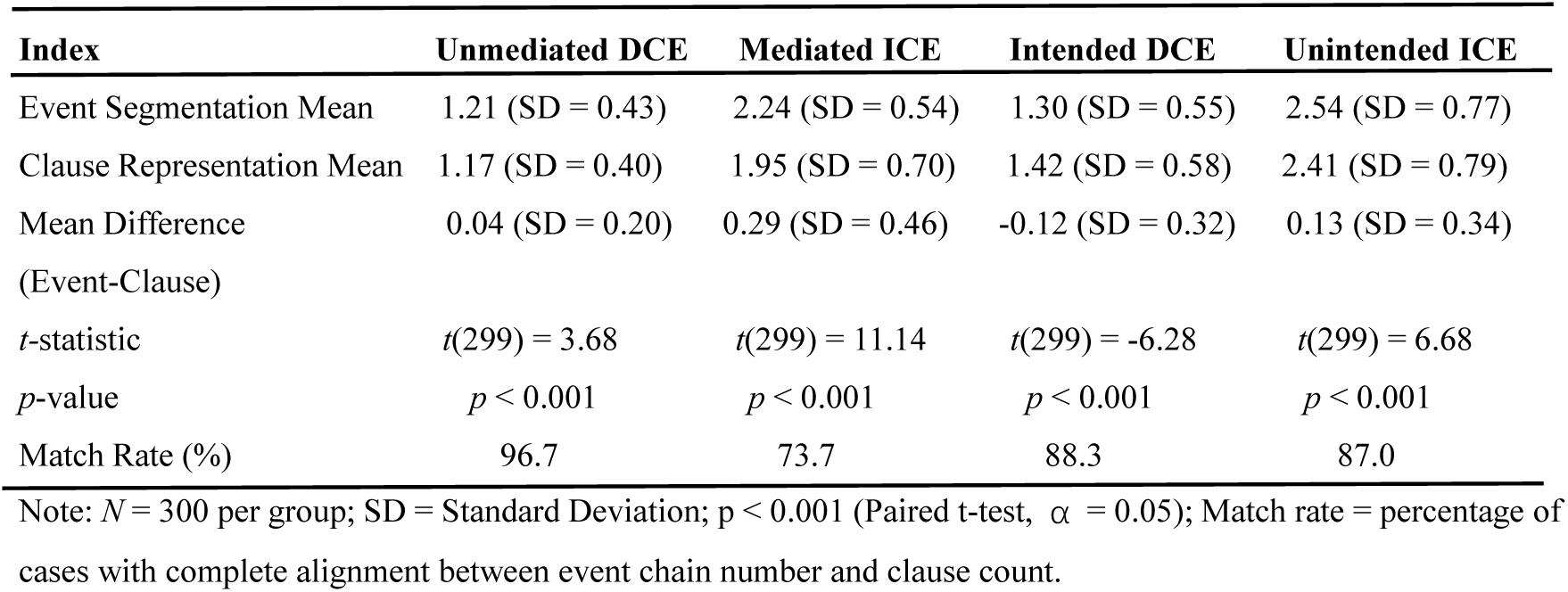
Paired *t*-test results: Event segmentation and clause representation in direct and indirect causal chains.

For indirect causal chains, mediated indirect causal events exhibited a shift toward multi-unit representations, with two-chain events (70%) and two-clause structures (55.3%) as the majority. The match rate (73.7%) was the lowest among the four types, with a significant difference (t (299) = 11.14, p < 0.001) showing more event units (M = 2.24, SD = 0.54) than clause units (M = 1.95, SD = 0.70). Unintended indirect causal events demonstrated the most complex representation pattern, with three-chain events (42.0%) and three-clause structures (33.6%) showing the highest percentages among all four types. Single-clause representation was the lowest (10.0%), while the use of three-clause (33.6%) and four-clause (8.7%) structures was the highest. The match rate was 87.0%, with a significant difference (t (299) = -6.68, p < 0.001) between event units (M = 2.54, SD = 0.77) and clause units (M = 2.41, SD = 0.79).

## 5. Discussion

### 5.1 Event segmentation and boundary assignment of direct/indirect causal chains

According to event segmentation theory, individuals naturally segment continuous experience into discrete (sub) events—bounded activity units with identifiable beginnings and ends (Zacks & Swallow 2007). This framework posits that a multimodal working event model is maintained during ongoing activities (Kaiser 2020), which persists as long as predictions are confirmed. Event boundaries occur when shifts in narrative dimensions—such as time, space, entities, causation, or intentionality—produce transient increases in prediction error, prompting model updating (Zacks 2004, 2020; Bailey et al. 2013). Our experimental findings provide robust empirical support for the construal distinction underlying the event segmentation of direct and indirect causal chains. For mediated and unmediated causal events, in the strong preference for single-chain segmentation unmediated direct causal events (79.7%), aligns with the theoretical expectation that the absence of intervening agency enables the conceptualization of the entire cause-effect sequence as an integrated event unit. Conversely, the introduction of an intervening agent in mediated indirect causal events fundamentally influences segmentation patterns. That is, observers demonstrate heightened sensitivity to event boundaries as the a secondary agent occurs, strategically adjusting the temporal grain, scope, and magnitude of their boundary reports in response. The significantly diversified segmentation patterns—with two-chain events predominating (70%)—aligns with the event-indexing model, wherein changes of protagonist trigger boundary detection.

As Langacker (1991: 294) notes, events afford multiple conceptual interpretations, and observers indeed adopt varying segmentation and boundary assignment strategies across causal scenarios, modulated by differing levels of granularity. Linguistic examples in which the number of segmented sub-events matched the number of clauses are selected as illustrative references for the interpretation. In unmediated direct causal events, coarse-grained segmentation establishes the event boundary at the achievement of the agent’s goal (e.g., “the cup broke”), resulting in single-chain events (Example 3a). Two-chain segmentation introduces sub-event boundaries at points where changes of protagonists or objects emerge, distinguishing, for instance, between the cause action of the agent (pushing the swing) and the goal effect on the patient (swing swinging) in Example 3b, or establishing boundary when the agent appears and isolating the patient’s state from the state-change event in Example 3c. Three-chain segmentation further decomposes the agent’s action goal into finer-grained actions and sub-goals (Example 3d), with boundaries set at the accomplishment of lower-level goals (e.g., “the egg broke” and “into the bowl”). Notably, the involvement of instrumental intermediaries in intended direct causal events often leads to a marked increase in two-chain segmentation (Example 3e). This may suggest that even inanimate instrumental intermediaries can trigger boundary assignment, underscoring the role of objects in shaping event models in Mandarin. Nevertheless, the predominance of single-chain event in this scenario suggests that inanimate participants lack intention and thus exert a less pronounced influence on event segmentation compared to animate participants.

(3) a. C4S10: 女孩砸碎了杯子。

Nǚhái zá suì le bēizi

Girl smash break ASP cup

‘The girl smashed the cup.’

b. C2S14: 一个男人推了一下秋千, 秋千就荡起来了。

yī gè nánrén tuī le yīxià qiūqiān qiūqiān jiù dàng qǐlai le

One-CL man push ASP one-CL swing swing then sway up come ASP

‘A man pushed the swing, and it started to sway.’

c. C8S14: 蜡烛燃着, 有一个人把蜡烛吹灭。

làzhú rán zhe yǒu yí gè rén bǎ làzhú chuī miè

Candle burn DUR exist one-CL person BA candle blow extinguish

‘The candle was burning, and a person blew it out.’

d. C5S14: 这个女生拿鸡蛋, 然后把鸡蛋打碎, 把鸡蛋放到碗里。

zhège nǚshēng ná jīdàn, ránhòu bǎ jīdàn dǎ suì, bǎ jīdàn fàng dào wǎn li

This-CL girl take egg then BA egg hit break, BA egg put arrive bowl inside

‘The girl took an egg, then broke it, and put it into a bowl.’

e. C10S11: 一个女生拿着锤子, 然后把盘子砸碎了。

Yī gè nǚshēng ná zhe chuízi ránhòu bǎ pánzi zá suì le

One-CL girl hold DUR hammer then BA plate smash break ASP

‘A girl took a hammer and then smashed the plate.’

In indirect causal events with intervening agents, coarse segmentation locates the event boundary at the primary agent’s goal achievement (Example 4a). In two-chain segmentation, more boundaries are set at the completion of the agent’s direct action and the initiation of the intermediary’s action (Example 4b). Finer-grained (three- and four-chain event) segmentation further partitions the actor’s and intermediary’s actions into discrete units bounded by lower-level action completion (Examples 4c-4d).

(4) a. C12S26: 一个女生让男生推秋千。

Yī gè nǚshēng ràng nánshēng tuī qiūqiān

One-CL girl let boy push swing

‘A girl asked a boy to push the swing.’

b. C14S18: 这个男士让另一男士把碗打碎, 然后他就把碗打碎了。

Zhège nánshì ràng lìng yī nánshì bǎ wǎn dǎ suì ránhòu tā jiù bǎ wǎn dǎ

This-CL man let another one-CL man BA bowl hit break then he then BA bowl hit

suì le

break ASP

‘The man asked another man to break the bowl, and then he broke it.’

c. C13S6: 这个女的让这个男的推一下秋千, 男的就推了一下秋千, 秋千就荡起了。

zhège nǚ de ràng zhège nán de tuī yīxià qiūqiān nán de jiù

This-CL woman let this-CL man push one-CL swing man then

tuī le yīxià qiūqiān qiūqiān jiù dàng qǐ le

push ASP one-CL swing swing then sway up ASP

‘The woman asked the man to give the swing a push. He pushed it, and the swing began to sway.’

d. C17S9: 红衣服女生示意黑衣服女生, 让她把台灯关掉, 然后黑衣服女生就照做了, 台灯灭了。

Hóng yīfu nǚshēng shìyì hēi yīfu nǚshēng ràng tā bǎ táidēng

Red clothes girl signal black clothes girl let her BA desk-lamp

guān diào ránhòu hēi yīfu nǚshēng jiù zhào zuò le táidēng miè le

turn off then black clothes girl then follow do ASP desk-lamp extinguish ASP

‘The girl in red signaled to the girl in black to turn off the desk lamp, which she then did, extinguishing the light.’

For intended & unintended causal chains, the parallel findings further substantiate the role of intentionality as a key factor in event segmentation. Goal-directed actions, which provide a coherent intentional framework, facilitated integration into single-chain events. In contrast, unintended causal events, lacking this unifying framework, were consistently parsed into two or more separate sub-events. This underscores that event segmentation is not solely driven by physical parameters but is profoundly shaped by conceptual factors such as causation and intentionality (Zwann & Radvansky 1995; Richmond et al. 2017), confirming that boundary assignment is a cognitively motivated process that serves to structure continuous experience. In unintended indirect causal events, native Mandarin speakers exhibit a strong preference for multi-chain segmentation (93%). Coarse-grained segmentation typically locates the event boundary at the completion of the causal sequence. In contrast, fine-grained segmentation introduces boundaries at points where event dimensions violate expectations, particularly with the appearance of a new entity or a goal-relevant change—prompting observers to separate the actor’s action endpoint from its outcome (Example 5a). This granularity can extend further: three-chain segmentation adds more boundaries at the completion of indirect causal sub-events, alongside the unexpected change of another entity (Example 5b), while four-chain segmentation further elaborates on protagonist actions or specific sub-results (Example 5c).

(5) a. C34S21: 一个人用铁锤砸了桌子, 但是把瓷碗给震碎了。

Yī gè rén yòng tiěchuí zá le zhuōzi, dànshì bǎ cíwǎn

One-CL person use iron-hammer smash ASP table but BA porcelain-bowl

gěi zhèn suì le

give shake break ASP

‘Someone smashed the table with an iron hammer, but ended up shattering the porcelain bowl.’

b. C31S2: 一个女的在弹球, 不小心球弹到桌子上, 把花瓶碰碎了。

Yī gè nǚ de zài tán qiú, bù xiǎoxīn qiú tán dào zhuōzi shàng

One-CL woman PROG bounce ball not carefully ball bounce arrive table top

bǎ huāpíng pèng suì le

BA vase bump break ASP

‘A woman was playing with a ball and accidentally bounced it onto the table, breaking the vase.’

c. C31S25: 一个女人弹着手上的球, 然后不小心球跳到了桌子上, 砸到了花瓶, 然后把花瓶砸碎了。

Yī gè nǚrén tàn zhe shǒu shàng de qiú, ránhòu bù xiǎoxīn

One-CL woman bounce DUR hand on DE ball then not carefully

qiú tiào dào le zhuōzi shàng, zá dào le huāpíng, ránhòu bǎ huāpíng

ball jump arrive ASP table top hit arrive ASP vase then BA vase

zá suì le

hit break ASP

‘A woman was bouncing the ball in her hand, and then accidentally the ball bounced onto the table, hitting the vase, and then she smashed the vase.’

A comparison across the four groups of causal chains reveals further insights into the role of intentionality and mediation in shaping event segmentation. Firstly, unintended indirect causal events exhibited the most fine-grained event segmentation among all conditions, with 41% of trials segmented as three-chain events and 10% as four-chain events. The lack of a clear intentional agent appears to prompt participants to parse the event into more fine-grained sub-units, reflecting a cognitive strategy of decomposing the causal sequence into smaller, more discrete action-outcome pairs. Secondly, mediated indirect causal events showed a different distribution, with two-chain segmentation (70%) being most frequent. This distinction highlights that the presence of an intervening agent prompts a moderate degree of segmentation — typically into two separate sub-events—whereas the absence of intentionality drives even finer-grained segmentation. The contrast may suggest that intentionality may serve as a particularly salient event boundary cue. This finding extends the event-indexing model (Zwaan & Radvansky 1998) in that among the five situational dimensions, intentionality may carry greater weight in segmentation than previously recognized, especially in causal event cognition.

Our study thus delineates the segmentation patterns of four types of causal chains, validating that prediction error facilitates boundary assignment and the establishment of new event models (Stawarczyk et al. 2019), while demonstrating that segmentation is influenced not only by temporal features (Bohnemeyer et al., 2010; Bohnemeyer & Van Valin, 2017). From the above analysis, it is evident that protagonists and objects, intentionality, endpoints of actions and causal relationships exert significant boundary effects, serving as key criteria for event boundary assignment. The findings demonstrate that the cognitive system goes beyond tracking movements and superficial features, instead prioritizing the understanding of actions, intentions, and objecthood. Goal processing enables observers to anticipate upcoming actions (Loucks & Pechey 2016). Coarse-grained segmentation anchors global boundaries at the achievement of the actor’s goals, whereas fine-grained segmentation dissects higher-level actions and goals, locating boundaries at the action endpoints, the introduction of new protagonists, the onset of new causal relations, or goal-relevant or unexpected change.

### 5.2 Semantic integration and specificity of direct/indirect causal chains

In accordance with the Theory of Event Coding (Hommel et al. 2001), event representations are composed of feature codes, among which those related to attractiveness and desirability are assigned greater weight (Zacks 2020). This weighting influences the specificity of event representations and ultimately affects their semantic integration. The cognitive processing of event coding involves two complementary stages: activation and integration. This view is consistent with Talmy’s theory of attention windowing (Talmy 2000). After continuous experience is segmented into discrete sub-events, the attention window functions as a selective mechanism (Fernandez-Duque & Johnson 1999; Lampert 2009), foregrounding specific sub-events within the conceptual complex through the cognitive operations of windowing and gapping (Talmy, 2000). Observers then integrate these foregrounded sub-events into conceptual chunks, which can be further assembled into higher-order event units, ultimately giving rise to a structured conceptual gestalt (Langacker 2008). Our experimental data provide empirical support for the role of specificity and semantic integration in the conceptual construction of causal chains.

At the most fundamental level, in unmediated direct causal chains, cause and effect not only display tight temporal alignment but also share the same patient, favoring integration into complex clauses — a pattern consistent with Givón’s (2001) proximity principle, whereby temporally co-occurring or spatially contiguous events are construed as a single gestalt. This packaging is realized through three primary integration patterns: (i) a macro-event “cause + effect” pattern encoded by verb-complement constructions (Example 6a); (ii) a “cause” pattern represented by a single verb, which profiles the agent’s action while omitting the effect event (Example 6b); and (iii) a “cause + effect” pattern encoded by *rang*-constructions (i.e., structures marked by the causative marker *rang*), which explicitly encode the agent, the effect event, and the causal relationship between them (Example 6c). This supports the theory of “metonymic clipping” (Van Valin & Wilkins 1996), whereby a conscious agent stands for the entire causal sequence. Notably, the introduction of an instrumental object triggers the use of a serial verb construction in Chinese. Despite the syntactic complexity, the entire event remains integrated into a single clause, demonstrating a conceptual merging of cause and effect that forms a coherent, higher-order event unit (Example 6d). The strong preference for highly integrated semantic packets (accounting for 56.7%) in direct causation, corroborates Givón’s (2001) principle that stronger semantic bonds yield tighter syntactic integration. This finding illustrates how tight conceptual packing enables syntactically condensed realizations, wherein cause and effect are conceptualized as a unified macro-event.

1. a. C4S28: 一个女孩砸碎了一个小碗。

Yí gè nǚhái zá suì le yí gè xiǎo wǎn

one-CL girl smash break ASP one-CL small bowl

‘A girl smashed a small bowl.’

b. C1S22: 她撕了一张纸。

Tā sī le yī zhāng zhǐ

She tear ASP one-CL paper

‘She tore a piece of paper.’

c. C2S8: 这位男士让秋千动了起来。

zhè wèi nánshì ràng qiūqiān dòng le qǐlai

this-CL man let swing move ASP up come

‘This man let the swing move.’

d. C10S6: 女子用锤子敲碎了盘子。

nǚzǐ yòng chuízi qiāo suì le pánzi.

woman use hammer hit break ASP plate

‘The woman smashed the plate with a hammer.’

In contrast, mediated indirect causal chains consistently prompt more detailed semantic integration. The primary agent’s control over the final outcome is reduced because animate intermediaries act consciously—this weakens the causal link between the initial action and end result. The intervening agent creates a cognitive boundary that requires fine-grained segmentation, producing a hierarchical conceptual structure where the primary agent’s action is embedded within the subsequent agent’s action. While periphrastic causative verbs allow limited single-clause compression of multi-agent events (Example 7a), the prevalence of coordinate clauses (Example 7b) and multi-clause structures reflects a conceptual need to split these causal chains into more discrete, sequentially organized units. This distributed encoding iconically mirrors the conceptual division of events into distinct agentive tiers. Frequent use of coordinate and extended conjunctive patterns reveals a sequential, relational assembly process: discrete sub-events are integrated into a complex sentence. The emergence of causative loop structures further confirms the cognitive complexity of representing mediated agency, which requires non-linear syntactic mappings (Example 7c). In this structure, two subordinate clauses are employed to emphasize the two actions of the intermediate agent, with both actions being generated from the sub-goal of the primary agent. Observers draw upon prior knowledge to structure events into a high-level partonomic hierarchy. Thus, fewer shared parameters between sub-events and weaker causal semantic associations make them more likely to be interpreted as multi-chain events, realized through less integrated clause structures.

Crucially, intentionality functions as a superordinate framing layer within semantic integration. Goal-directed actions, organized around a unified intentional schema, favor integrated packaging—even within single clauses—suggesting that intentionality provides a conceptual scaffold that supports semantic integration. In contrast, unintended causal event triggers markedly different packaging strategies: the use of adversative conjunctions and explicit markers of accidentality reveals a conceptual need to explain unexpected outcomes, resulting in more elaborated, often multi-clausal descriptions that segment the sequence into a causal event plus mitigating circumstances (Example 7d). Lacking a cohesive intentional frame, unintended events are constructed as looser sub-event assemblages, requiring explicit unexpectedness markers to reconcile causal discrepancies. This demonstrates that the conceptual hierarchy is not linear but interactive, with intentional framing modulating integration strength between segmented components.

(7) a. C12S10: 一个人让他晃动秋千。

yí gè rén ràng tā huàng dòng qiūqiān.

one-CL person let him swing move swing

‘A person let him swing the swing.’

b. C19S29: 女一号让女二号把纸剪了, 然后女二号就把纸剪了。

nǚ yī hào ràng nǚ èr hào bǎ zhǐ jiǎn le, rán hòu

woman one number let woman two number BA paper cut ASP then

nǚ èr hào jiù bǎ zhǐ jiǎn le.

woman two number then BA paper cut ASP

‘The first woman lead made the second woman cut the paper, and then the second woman cut the paper.’

c. C19S4: 黑衣服女生按照粉衣服女生的要求, 把蜡烛吹灭了。

hēi yīfu nǚshēng ànzhào fěn yīfu nǚshēng de yāoqiú,

black clothes girl follow pink clothes girl DE request

bǎ làzhú chuī miè le

BA candle blow extinguish ASP

‘The girl in black clothes blew out the candle according to the request of the girl in pink clothes.’

d. C31S2: 一个女的在弹球, 不小心球弹到桌子上, 把花瓶碰碎了。

Yī gè nǚ de zài tán qiú, bù xiǎoxīn qiú tán dào zhuōzi shàng

One-CL woman PROG bounce ball not carefully ball bounce arrive table top

bǎ huāpíng pèng suì le

BA vase bounce break ASP

‘A woman was playing with a ball and accidentally bounced it onto the table, breaking the vase.’

Together, these findings affirm that event coding reflects not merely informational detail, but the degree of conceptual specificity and semantic integration. As Talmy (2000: 276) observes, the cognitive representation of causal chains exhibits functional stability while permitting contextually adaptive, plastic representational forms—a phenomenon particularly evident in high-specificity expressions. This perspective is corroborated by Langacker (1987: 345), who states that cognitive grammar sanctions all patterns with varying degrees of productivity and regularity, encouraging the expectation that causative structures may assume different forms as parameters vary. These results collectively reveal a systematic hierarchy of semantic integration underlying the conceptual construction of causal chains, where direct and indirect causal events occupy distinct positions along a specificity continuum. The semantic integration and clausal union observed across four types of causal scenarios validates the role of eventhood and agentivity, which is governed not only by co-temporality and spatial contiguity but also by intentionality, co-reference, coercive power and control (Givón 2001, 2009). Furthermore, direct causal events exhibit high stability in conscious representation, while indirect causal events are characterized by lower predictability and greater conceptual plasticity.

### 5.3 Conceptual construction hierarchy of direct/indirect causal chains

Cross-linguistic evidence confirms that direct and indirect causal chains represent universal semantic categories in causal conceptualization (Wolff 2007; Bohnemeyer et al. 2007, 2011). English typically encodes this distinction through lexical versus periphrastic causatives (e.g., “Sarah opened the door” vs. “Sarah caused the door to open”). Chinese, however, exhibits richer variation in linguistic coding of these causation types, revealing deeper cognitive-semantic regularities. The analysis of the relationship between cognitive segmentation and clausal representation revealed three main findings. First, direct causal events exhibited high match rates and a predominance of single-chain segmentation. This pattern aligns with the event-indexing model (Zwaan & Radvansky 1998) and suggests that direct causal events form coherent conceptual units that are naturally encoded as single clauses. Second, a key discovery lies in the distinct pattern observed for unintended indirect causal events, which showed significantly higher rates of three-chain segmentation and three-clause representation compared to mediated indirect causal events. This indicates that intentionality functions as a more powerful segmentation breakpoint than mediation in guiding event cognition. Two mechanisms may explain this phenomenon. On the one hand, unintended causal events lack agentive control, thereby creating clearer conceptual boundaries between sub-events, which speakers perceive as distinct units. On the other hand, the absence of intentionality necessitates more explicit linguistic marking of event boundaries, leading to a higher use of multi-clause structures. While shifts in protagonists and causation have been highlighted as particularly salient for segmentation (McNerney et al. 2011), the current findings extend previous research by demonstrating that intentionality not only affects segmentation salience but also modulates the mapping between cognitive segmentation and linguistic representation. Third, indirect causal events exhibit lower match rates and more variable unit counts, reflecting flexible mapping between event segmentation and clause representation. The significant *t*-test results (*p* < 0.001), which show more event units than clause units in both mediated and unintended causal types, indicate active semantic integration processes: speakers frequently merge multiple segmented sub-events into fewer clauses when causal relationships are indirect. This flexibility is particularly evident in mediated indirect causation, suggesting that mediation creates optional integration points rather than mandatory boundaries.

Based on these findings, we propose an integrated cognitive-linguistic model, conceptually outlined in Figure 7, which delineates the pathway from event perception to syntactic representation. This process segments the entire cause-effect sequence into cognitively atomic units and integrates these sub-events, driving a bottom-up construction that progresses from a unified conceptual core to syntactically compact forms. Anchored in continuous experience, the model identifies event segmentation as the foundational cognitive operation for constructing causal chains. This segmentation operates at either coarse or fine granularity, with the chosen level directly governing the hierarchy of conceptual construction. Coarse-grained segmentation, typically applied to highly unified events such as direct causation, leads to low specificity in syntactic representation, yielding tight clausal union and compact syntactic structures such as verb-resultative compounds. Conversely, under fine-grained segmentation, guided by the windowing of attention mechanism, event components acquire varying degrees of salience. Observers foreground certain sub-events through attentional windowing, leading to semantic integration at differing levels of specificity. The granularity of semantic integration and clausal union are modulated by core cognitive factors such as eventhood and agentivity. For instance, indirect causal events involving an intermediate agent or non-goal-directed actions exhibit reduced semantic integration, which manifests as loosely connected, multi-clausal structures. This cognitive interplay ultimately crystallizes into linguistic expressions of varying syntactic tightness.

**Fig 7.**
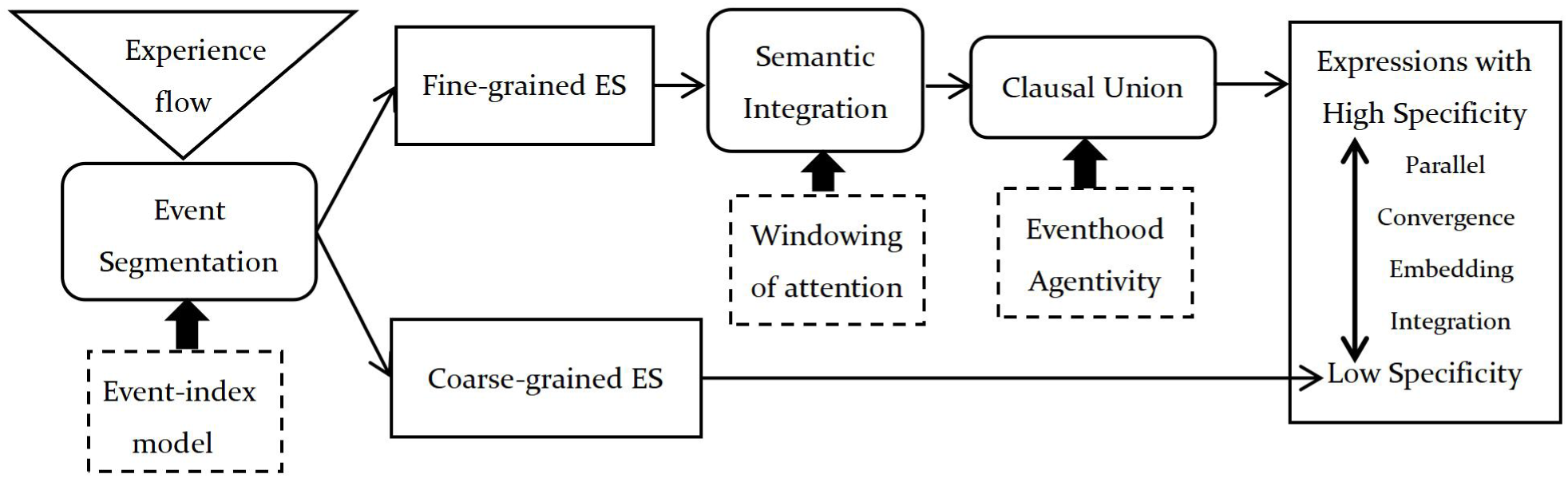
Conception Construction Model of Causal Chains (based on Zacks 2020; Talmy 2000 & Givón 2009)

The model posits that this cognitive-syntactic correspondence is governed by several principal mechanisms. First, the granularity of conceptual segmentation guided by event-indexing model modulates linguistic specificity at syntactic level. Coarse-grained segmentation condenses causal sequences into single event units, typically encoded as single-clause expressions. This segmentation strategy fosters the emergence of concise structural gestalts and motivates the embedding and fusion of adjacent syntactic constituents, thereby aligning with the economy principle of structural organization. On the contrary, fine-grained segmentation parses events into multiple conceptual sub-events, which tend to be encoded through more expansive syntactic configurations such as causal compound sentences or clause clusters.

Second, semantic integration shapes the syntactic coding of clause-structural gestalts. In perceiving causal events, native Chinese speakers are inclined to track actions, intentions and protagonists rather than mere spatiotemporal features. The fulfillment of the actor’s goal and the completion of goal-relevant actions attract heightened attention and engage language-processing mechanisms that support chained embedding and conjunction. This gives rise to single-clause expressions such as simple verb-predicate sentences, compact complex constructions (e.g., verb-complement and pivotal structures), and embedded complex sentences (e.g., double-object predicates). Furthermore, packaging sub-events at junctures where new protagonists enter or new causal relations are introduced fosters the development of inter-clausal syntactic relations—such as coordination and conjunction — resulting in two-chain integration patterns including causal succession, coordination, bifurcation, and looping. More specific cognitive construction dissects and elaborates the goals or sub-goal and segmentation of protagonists, encoding causal events through more loosely organized, parallel multi-clause structures that yield mixed or extended causal clause clusters.

This conceptual construction hierarchy offers a unified explanatory framework for the mind-language interface in causation: the conceptual organization of a causal event — is not a direct reflection of the world but a cognitive process of the segmentation of experience, the semantic integration of sub-events and the syntactic devices of a language, influenced by the salient parameters regarding agency, intentionality and causal relationship. The syntactic representation of causation, as a form-meaning pairing, is iconically mapped onto a continuum of clause integration—from loosely connected to tightly integrated syntactic constructions. High-specificity expressions stem directly from fine-grained segmentation and low integration, employing syntactic strategies such as embedding to represent hierarchical complexity, along with parallel or coordinate structures to elaborate discrete sub-events. In contrast, low-specificity expressions reflect coarse-grained segmentation and high integration, favoring monolithic, convergent syntactic packaging that presents complex scenes as unified conceptual gestalts.

## 6. Conclusion

The conceptual construction of causal chains generates the organization continuum of clause structures ranging from loose to integrated. The syntactic representation at the linguistic level mirrors the segmentation and integration of events at the conceptual level. While the iconic mapping between semantics and syntax is widely acknowledged, the specific cognitive and conceptual mechanisms that govern this isomorphism require further articulation. Motivated by this gap, this study investigates how linguistic representations of direct and indirect causal events in Chinese vary, and elucidates how these variations are systematically rooted in cognitive segmentation and integration, by comparing experimental data across four causal scenarios.

Based on foundational insights from cognitive and linguistic research, we propose a conceptual construction model for the event coding of causal chains. Anchored in a coherent theoretical triad—comprising event segmentation, semantic integration, and clause union — this model offers an integrated framework for investigating the mind-language interface. In causal events, segmentation granularity varies and is modulated by an event-indexing mechanism. Following segmentation, the resulting sub-events undergo attentional windowing and semantic integration, processes that underlie the cognitive structuring and specificity of event representation. These integrated conceptual structures are then iconically mapped onto syntactic organization, systematically encoded as variations in clausal architecture and linguistic form.

Our findings consistently reveal that Chinese native speakers track intentions, protagonists and accomplishment of actions in causal event segmentation. On one hand, corresponding with the linguistic tendency, Chinese speakers—like German speakers—focus more on action endpoints than Arabic speakers do (Flecken et al. 2014). Chinese speakers assign considerable attention to the achievement of goals or sub-goals and goal-relevant actions. Event models are rapidly established or updated through prediction errors along these dimensions (Loucks & Pechey 2016). Coarse-grained segmentation anchors global boundaries at the attainment of the actor’s goals, whereas fine-grained segmentation parses higher-level actions and goals by locating boundaries at action endpoints, the introduction of new protagonists, the onset of new causal relations, or goal-relevant or unexpected changes. On the other hand, as a language that can use some constructions to package a sequence of smaller actions into a single event, Chinese allows speakers to span the entire verb string—as seen in serial verb constructions and periphrastic causative structures—suggesting that they conceptualize it as an integrated unit.

These results provide robust empirical support for the principle of syntactic iconicity (Givón 2001; Haiman 1983), demonstrating that the “strength of the semantic bond” is not merely an abstract linguistic notion but a predictable outcome of cognitive segmentation and conceptual integration. We have thus advanced from description to explanation by unraveling the operation of the mind-language interface in the domain of causation: the granularity of cognitive segmentation governs semantic integration, which in turn is reflected in syntactic union and linguistic specificity. This work has twofold implications for relevant theories. First, it bridges cognitive and linguistic accounts of causation by proposing a unified model that explains linguistic patterning through cognitive operations. By demonstrating that intentionality and mediation differentially affect the mapping between event segmentation and linguistic representation, it extends the event-indexing model (Zwaan & Radvansky 1998) in specifying how cognitive dimensions interact. It identifies the conceptual dimensions to which Chinese speakers accord heightened attentional weighting in causal event segmentation. We suggest that intentionality, protagonists, and accomplishment of action are critical boundary cues in causal events segmentation. Critically, these conceptual dimensions systematically modulate segmentation granularity and semantic integration — even when spatio-temporal cues exhibit minimal variation. Second, evidence from cases of fine-grained segmentation with integrated syntactic structures and variable segmentation-representation mapping in indirect causal events confirms semantic integration and clause union in linguistic production processes and demonstrates that clause union is sensitive to syntactic structures in specific language. It elucidates the diversity in Chinese causal representation and clarifies the underlying conceptual schemas and cognitive mechanisms. These patterns suggest universal cognitive tendencies while highlighting language-specific strategies. Several limitations of the present study should be acknowledged. First, the non-verbal segmentation task relied on verbal instructions involving the terms “(sub)event,” which may have introduced mild linguistic construal bias despite the use of short, simple causative video stimuli. Second, the study focused on offline behavioral responses rather than online cognitive processes, so the precise temporal dynamics underlying the integration of these cues remain to be further investigated. Future research could extend this paradigm to other linguistic domains and typologically diverse languages to test the universality of this construction hierarchy. Furthermore, employing neuroimaging techniques could illuminate the real-time neural correlates of the segmentation and integration processes identified here, offering a yet deeper glimpse into the cognitive architecture of causation.

## Notes

### Competing Interest Statement

The authors have declared no competing interest.

## References

Bailey, H. R., Kurby, C. A., Giovannetti, T, & Zacks, J. M. (2013). Action perception predicts action Performance. Neuropsychologia, 51, 2294–304.

Bohnemeyer, J., Enfield N. J., Essegbey J., & Kita S. (2010). The macro-event property: The event segmentation of causal chains. In Bohnemeyer, J. & E. Pederson (eds.). Event Representation in Language and Cognition (pp.44–67). Cambridge: Cambridge University Press.

Bohnemeyer, J., Enfield, N., Essegbey, J., Ibarretxe, I., Kita, S., Lüpke, F., & Ameka, F. K. (2007). Principles of event representation in language: The case of motion events. Language, 83(3): 495–532.

Bohnemeyer, J. & Van Valin, R. D. (2017). The macro-event property and the layered structure of the clause. Studies in Language, 41(1):142–197.

Davidson, D. (1969). The individuation of events. In Rescher N.(ed.) Essays in Honor of Carl G. Hempel (pp. 216–234). Dordrecht: Reidel.

Deng, Y., & Li, T. F. (2023). Event segmentation and causation: the cause of Mandarin causal-chain motion. Studia Linguistica, 77(2), 368–415.

Fernandez-Duque, D. & L. J. Mark. (1999). Attention metaphors: How metaphors guide the cognitive psychology of attention. Cognitive Science, 23(1), 83–116.

Flecken, M., von Stutterheim, C., & Carroll, M. (2014). Grammatical aspect influences motion event perception: Evidence from a cross-linguistic non-verbal recognition task. Language and Cognition, 6(1), 45–78.

Gerwien, J. & von Stutterheim, C. (2018). Event segmentation: Cross-linguistic differences in verbal and non-verbal tasks. Cognition, 180, 225–237.

Givón, T. (1991). Serial verbs and the mental reality of ‘event’. In Traugott E. C. & B. Heine (eds.), Approaches to grammaticalization,vol.1(pp.81–127). Amsterdam:John Benjamins.

Givón, T. (2001). Syntax: An Introduction, Vol. II, Amsterdam: John Benjamins.

Givón T. (2009). The Genesis of Syntactic Complexity: Diachrony, Ontogeny, Neuro-cognition, Evolution[M]. John Benjamins Publishing.

Haiman, J. (1983). On some origins of switch-reference marking, In Haiman J. & P. Munro (eds). Switch Reference and Universal Grammar, TSL #2, Amsterdam: John Benjamins.

Hommel, B., Muesseler, J., Aschersleben, G., & Prinz, W. (2001). The Theory of Event Coding (TEC): a framework for perception and action planning. Behav. Brain Sci., 24, 849–937.

Kaiser, E. (2020). Linguistic consequences of event segmentation in visual narratives: implications for prominence. Language, Cognition and Neuroscience, 35(3), 402–408.

Keven, N. (2016). Events, narratives and memory. Synthese, 193, 2497–2517.

Krebs, J., Harbour, E., Malaia, E. A., Wilbur, R. B., Martetschläger, J., Schwameder, H., Roehm, D. (2025). Sign language encodes event structure through neuromotor dynamics: motion, muscle, and meaning. Frontiers in Psychology (Preprint).

Lampert, M. (2009). Attention and Recombinance. Frankfurt am Main: Peter Lang Press.

Langacker, R. (1987). Foundations of Cognitive Grammar. Vol. 1: Theoretical Prerequisites. Standford: Stanford University Press.

Langacker, R. W. (1991). Foundations of Cognitive Grammar, Vol. II: Descriptive Application. Stanford: Stanford University Press.

Langacker, R. W. (2008). Cognitive Grammar: A Basic Introduction. Oxford: Oxford University Press.

Liu, C. (2016). Boundary assignment: The cognitive motivation of lexicalization. Modern Foreign Languages, 4, 449–458.

Loucks, J., & Pechey, M. (2016). Human action perception is consistent, flexible, and orientation dependent. Perception, 45,1222–1139.

McNerney, M. W., Goodwin, K. A., & Radvansky, G. A. (2011). A novel study: A situation model analysis of reading times. Discourse Processes, 48, 453–474.

Newtson, D. (1973). Attribution and the unit of perception of ongoing behavior. Journal of Personality and Social Psychology, 28, 28–38.

Papafragou, A. & Grigoroglou, M. (2019). The role of conceptualization during language production: evidence from event encoding. Language, Cognition and Neuroscience, 34(9), 1117–1128

Pawley, A. (1987). Encoding events in Kalam and English: different logics for reporting experience, In Tomlin R. S. (ed.), Coherence and Grounding in Discourse(pp.329–360). Amsterdam: Benjamins.

Radvansky, G. A., & Zacks, J. M. (2014). Event Cognition. Oxford: Oxford University Press.

Radvansky, G. A. (2017). Event Segmentation as a Working Memory Process. Journal of Applied Research in Memory and Cognition, 6, 121–123.

Richmond, L. L., Gold, D. A., & Zacks, J. M. (2017). Event perception: translations and applications. Journal of Applied Research in Memory and Cognition, 6(2),111–120.

Slobin, D. I. (1996). From “thought and language” to “thinking for speaking”. In Gumperz J. J. & S. C. Levinson (eds.), Rethinking Linguistic Relativity (pp.70–96). Cambridge: Cambridge University Press.

Stawarczyk, D., Bezdek, M. A., & Zacks, J. M. (2019). Event representations and predictive processing: the role of the midline default network core. Topics Cognitive Science, 13(1), 164–186.

Talmy, L. (2000). Toward a Cognitive Semantics. Volume I: Concept Structuring Systems. Cambridge, MA: MIT Press.

Van Staden, M. & Narasimhan, B. (2013). Granularity in the cross-linguistic encoding of motion and location. In Vulchanova M. & E. Van der Zee (eds.), Motion encoding in language and space(pp. 134–148). Oxford: Oxford University Press.

Van Valin, R. D., & Wilkins, D. P. 1996. The case for ‘effector’: case roles, agents and agency revisited. In Shibatani M. & S. Thompson (eds.), Grammatical Constructions: Their Form and Meaning (pp. 289–322). Oxford: Oxford University Press.

Wang, Y. C., Adcock, R. A., & Egner, T. (2023). Toward an integrative account of internal and external determinants of event segmentation. Psychonomic Bulletin & Review, 31, 484–506.

Wolff, P. (2003). Direct causation in the linguistic coding and individuation of causal events. Cognition, 88, 1–48.

Wolff, P, Jeon, G., & Li, Y. (2009). Causers in English, Korean, and Chinese and the individuation of events. Language and Cognition, 1(2),167–196.

Zacks, J. M. (2004). Using movement and intentions to understand simple events. Cognitive Science, 28, 979–1008.

Zacks, J. M. (2020). Event perception and memory. Annual Review of Psychology, 71(1), 165–191.

Zacks, J. M., & Swallow, K. M. (2007). Event segmentation. Current Directions in Psychological Science, 16(2), 80–84.

Zacks, J. M., & Tversky, B. (2001). Event structure in perception and conception. Psychological Bulletin, 127(1), 3–21.

Zacks, J. M., & Tversky, B. (2013). Granularity in taxonomy, time, and space. In Vulchanova M. & E. van der Zee(eds.), Motion encoding in language and space(pp.123–133). Oxford: Oxford University Press

Zwaan, R. A. (2016). Situation models, mental simulations, and abstract concepts in discourse comprehension. Psychonomic Bulletin & Review, 23,1028–1034.

Zwaan, R. A., & Radvansky, G. A. (1998). Situation models in language comprehension and memory. Psychological Bulletin, 123(2), 162–185.

